# Association between long-term air pollution exposure and DNA methylation: the REGICOR study

**DOI:** 10.1101/404483

**Authors:** Sergi Sayols-Baixeras, Alba Fernández-Sanlés, Albert Prats, Isaac Subirana, Michelle Plusquin, Nino Künzli, Jaume Marrugat, Xavier Basagaña, Roberto Elosua

**Author notes:** **Author for correspondence:** Roberto Elosua, MD, PhD, IMIM, Hospital del Mar Medical Research Institute, Dr Aiguader 88, 08003 Barcelona, Catalonia, Spain.

## Abstract

**Background:** Limited evidence suggests that epigenetic mechanisms may partially mediate the adverse effects of air pollution on health. Our aims were to identify new genomic loci showing differential DNA methylation associated with long-term exposure to air pollution and to replicate loci previously identified in other studies.

**Methods:** A two-stage epigenome-wide association study was designed: 630 individuals from the REGICOR study were included in the discovery and 454 participants of the EPIC-Italy study in the validation stage. DNA methylation was assessed using the Infinium HumanMethylation450 BeadChip. NOX, NO2, PM10, PM2.5, PMcoarse, traffic intensity and traffic load exposure were measured according to the ESCAPE protocol. A systematic review was undertaken to identify those cytosine-phosphate-guanine (CpGs) associated with air pollution in previous studies and we screened for them in the discovery study.

**Results:** In the discovery stage of the epigenome-wide association study, 81 unique CpGs were associated with air pollution (p-value <10^−5^) but none of them were validated in the replication sample. Furthemore, we identified 12 CpGs in the systematic review showing differential methylation with a p-value fulfilling the Bonferroni criteria and 1642 CpGs fulfilling the false discovery rate criteria, all of which were related to PM_2.5_ or NO_2_. None of them was replicated in the discovery study, in which the top hits were located in an intergenic region on chromosome 1 (cg10893043, p-value=6.79·10^−5^) and in the *PXK* and *ARSA* genes (cg16560256, p-value=2.23·10^−04^; cg11953250, p-value=3.64·10^−04^).

**Conclusions:** Neither new genomic loci associated with long-term air pollution were identified, nor previously identified loci were replicated. Continued efforts to test this potential association are warranted.

## BACKGROUND

Exposure to air pollution remains a global threat with more than 90% of the world’s population now exceeding the exposure limits proposed for particulate matter by the World Health Organization (WHO) (1). At the same time, a growing body of evidence consistently supports the adverse health effects of air pollution, which the same WHO report estimates to be related with 3 million premature deaths worldwide each year. However, the mechanisms by which air pollution induces these deleterious effects are not completely understood.

Epigenetics encompasses mechanisms that regulate gene expression without changing the DNA sequence, and may contribute to the relation between air pollution and health. The most studied epigenetic mechanism is DNA methylation, which is heritable but can also be modified by life-style and environmental factors. Recently, several studies have analyzed the association between air pollution and DNA methylation using a genome-wide approach, and have reported numerous loci showing differential methylation related to this exposure (2–10). The aims of this study were both to identify new genomic loci showing differential methylation associated with long-term exposure to air pollution in a population-based study in Spain and to replicate loci previously reported in other studies.

## METHODS

### Identification of new genomic loci showing differential methylation related to long-term air pollution exposure (Aim 1)

#### Study design and population

We designed a cross-sectional epigenome-wide association study in two stages. We used the REGICOR (REgistre GIroní del COR) cohort as the discovery study and the Italy center of the European Prospective Investigation into Cancer and Nutrition (EPIC-Italy) as the replication study, followed by a meta-analysis of the results observed in both studies (REGICOR + EPIC-Italy).

As previously described,(11) the REGICOR discovery sample included 648 participants randomly selected from the second wave of the REGICOR study in 2008-2013. The initial survey, performed during 2003-2005, included participants aged between 35 and 79 years, not institutionalized, and residing in Girona province (Catalonia, Spain) (11).

The EPIC-Italy replication study included 47,749 individuals in a multicenter prospective cohort recruited during 1993-1998 (12,13). The samples selected for the present study were from two case-control studies on breast cancer (14) and colorectal cancer in Varese and Turin.

#### HumanMethylation450 BeadChip

DNA was extracted using standardized methods from peripheral blood (Puregen TM; Gentra Systems) and buffy coats (QIAGEN QIAsymphony DNA Midi Kit) in the REGICOR and EPIC-Italy studies, respectively. DNA was bisulphite-converted and the epigenome-wide methylation profiles were obtained using the Infinium HumanMethylation450 BeadChip (Illumina) (450K) to assess methylation on 485,577 cytosine-phosphate-guanine (CpGs) throughout the genome, following the Illumina Infinium HD Methylation protocol (15,16). The REGICOR samples were processed in two centers of the Spanish National Genotyping Center: the Center for Genomic Regulation in Barcelona (n=188 samples) and the Centro Nacional de Investigaciones Oncológicas in Madrid (n=460 samples). All the processed batches contained two duplicate samples used as an internal quality control. The EPIC-Italy samples were analyzed at the Human Genetics Foundation in Turin. The same well-defined pipeline was used in both studies to assess the quality control of the methylation data (17). We used the M-value as the main DNA methylation measurement (Equation 1). An M-value=0 indicates that the CpG is half methylated, a positive M-value that the CpG is more methylated than unmethylated, and a negative M-value the inverse result. We standardized the M-value by batch (Equation 2) to reduce the batch effect and other potential technical sources of variation.

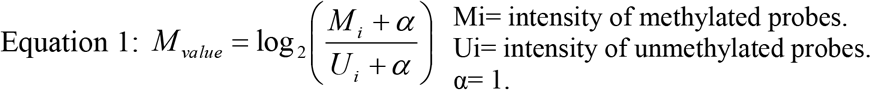

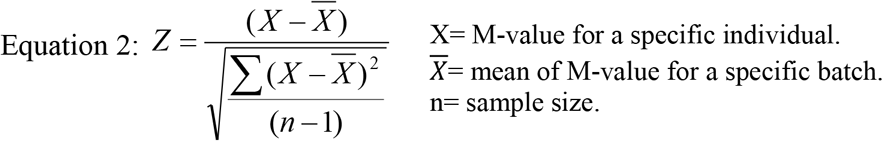

#### Air pollution exposure

Both the REGICOR data and the Turin component of EPIC-Italy contained particulate matter exposure [aerodynamic diameter of <10μm (PM_10_), <2.5μm (PM_2.5_), and PM_coarse_ (the difference between PM_10_ and PM_2.5_)], nitrogen oxides (NO_x)_ and nitrogen dioxide (NO_2_) measurements. For the Varese component of EPIC-Italy, only NO_x_ and NO_2_ were available. Both studies used the ESCAPE protocol to assess long-term exposure to air pollution (18,19). As previously described, address histories for the past 10 years were collected by questionnaire, and each address was geocoded at the front-door level (20). Using land use regression (LUR) models, 10-year weighted average exposure to NO_X_ and to nitrogen dioxide NO_2_ for each participant were estimated (21). The model’s coefficient of determination for NO_2_ was 0.63 for REGICOR and 70% for Turin, and for NO_x_ was 66% and 72%, respectively (20).

PM_10_, PM_2.5_ and PM_coarse_ were also assessed using LUR models. The R^2^ of the models for REGICOR and Turin was 71% and 69% for PM_10_, 51% and 59% for PM_2.5,_ and 71% and 58% for PM_coarse_, respectively (19).

We also used traffic proximity markers as surrogates of air pollution exposure in independent analyses. For each address, we calculated the traffic intensity at the nearest street and the traffic load (sum of traffic intensity multiplied by length of road segment) for all segments in a 100 meters buffer and derived 10-year average values for each participant.

#### Other covariates

REGICOR’s trained team of nurses collected relevant sociodemographic, lifestyle, and cardiovascular risk factors using standardized and validated questionnaires (22,23). Smoking exposure was grouped in four categories: current smoker (smoked ≥1 cigarette/day at the time of the visit, on average, or gave up smoking within the year of the visit); former smoker, 1-5 years (gave up smoking up to 5 years before the visit); former smokers >5 years; and never smokers. The EPIC-Italy study collected the same variables, as well as the participating center and patient’s diagnostic status.

We estimated cell concentration using Houseman algorithm by means of *minfi* R package (24,25). In both cohorts, we also calculated surrogate variables to control for potential residual cofounding, using the *sva* R package (26). These variables identify and remove potential and non-measured sources of variation due to technical and biological confounders.

#### Statistical analysis

We assessed the association between air pollution and DNA methylation using robust linear regression model to reduce the effect of outliers. We used the differing air pollution exposures as independent variables and DNA methylation as the dependent variable. The models were adjusted for age, sex, smoking exposure, and cell composition (Model 1). Moreover, a second model adjusted for surrogate variables was also fitted (Model 2). In the REGICOR discovery study, we selected for validation those CpGs with a p-value of the association below an arbitrary threshold (p-value <10^−5^) for each specific exposure. The results were replicated in the EPIC-Italy study using the same method and models, including also disease status and study center as additional covariates. The results for each CpG were then meta-analyzed using a random effects model, applying Bonferroni criteria to declare a result as statistically significant (0.05/427,948 CpGs; p-value<1.17·10^−07^).

### Replication of previously published CpGs associated with air pollution (Aim 2)

We performed a systematic review to identify relevant epigenome-wide association studies indexed in Pubmed (https://www.ncbi.nlm.nih.gov/pubmed/) from its inception to March 2018. We used the following search terms strategy: (methylati* [Title/Abstract]) AND (epigenome-wide [Title/Abstract] OR genome-wide [Title/Abstract] OR 450K [Title/Abstract] OR 450 [Title/Abstract] OR HumanMethylation450 [Title/Abstract]) AND (“Air pollution” [Title/Abstract]). The articles identified were manually screened by 1 reviewer (SS-B), focusing first on the title and abstract and then on the complete manuscripts to assess their appropriateness for inclusion in the review. The same author extracted the CpGs that were significantly associated with air pollution and replicated in at least one external population. In case of doubts the article was evaluated by a second reviewer (RE) to achieve a consensus. The identified CpGs were then screened in the selected REGICOR cohort.

#### Statistical analysis

The same analysis strategy and methodology described above was followed for the replication. We selected as distinctive the CpGs reported as statistically significant in the original studies, based on both Bonferroni corrected p-values and false discovery rate (FDR) p-values<0.05. We considered as replicated those CpGs that fulfilled both the Bonferroni criteria according to the number of CpGs previously discovered and the FDR p-value.

## RESULTS

### Identification of new genomic loci showing differential methylation related to air pollution (Aim 1)

#### Discovery Stage

After applying the quality control of the 450K array, we excluded 3 individuals and 57,629 CpG probes. Moreover, we removed those individuals without information on air pollution exposure (n=15). Finally, 630 individuals and 427,948 probes were included in the analysis. The main sociodemographic and clinical characteristics and the air pollution exposures of the study participants are shown in Table 1. The Manhattan plots and q-q plots of the associations between air pollution exposures and DNA methylation are shown in Supplementary Figure 1. We identified 81 unique CpGs associated with air pollution exposures with a p-value<10^−5^. In model 1, 6 CpGs were associated with PM_10_, 0 related to PM_2.5_, 5 to PM_coarse_, 2 to NO_X_, 4 to NO_2,_ and 2 to traffic at the nearest street (Supplementary Table 1). In model 2, 7 CpGs were associated with PM_10_, 18 related to PM_2.5_, 6 to PM_coarse_, 28 to NO_X_, 32 to NO_2,_ and 9 to traffic at the nearest street (Supplementary Table 2).

**Table 1:**
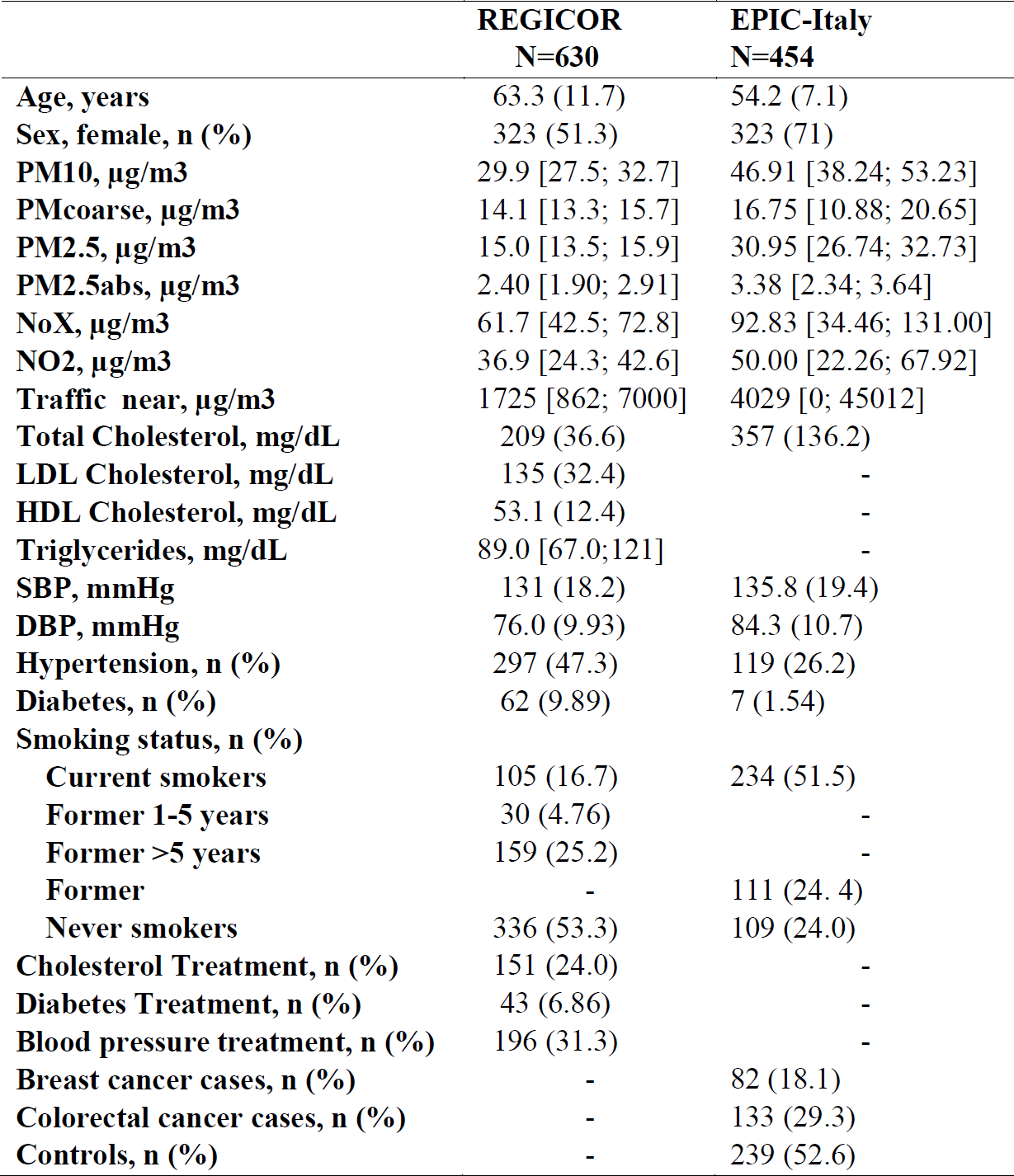
Main characteristics of the participants included in the discovery and validation studies.

#### Validation Stage and Meta-analysis

After applying a similar quality control of the 450K array, we included all 81 CpGs selected for replication and we excluded 5 individuals. Moreover, we removed those individuals without information on confounder variables (n=15) and air pollution exposure (n=61). Finally, 454 individuals were included in the analysis of NO traits (Turin and Varese) and 297 in the analysis of PM traits (Turin). The main characteristics and air pollution exposures of the EPIC-Italy participants are shown in Table 1. The associations between air pollution and the selected CpGs in this analysis are shown in Supplementary Table 1 and 2.

The results of the meta-analysis of the REGICOR and the EPIC-Italy studies are shown in Supplementary Table 1 and 2. None of the selected CpGs was validated in the joint analysis.

#### Statistical power

The regression coefficient values (effect size) that we could detect as statistically significant in the meta-analysis, accepting an alpha risk of 1.17·10^−07^ in a two-sided test and with an 80% power, are shown in Supplementary Table 1 and 2.

### Replication of the previously published CpGs associated with air pollution (Aim 2)

We initially identified 19 manuscripts (2–10,27–36) based on the search terms and after reading the full manuscripts we finally selected 9 studies (2–10). We selected 12 CpGs showing differential methylation in relation to PM_2.5_ from the only manuscript that corrected for Bonferroni criteria (9). Three of these CpGs could not be analyzed in the REGICOR study as they did not pass the quality control, and none of the others was replicated (Supplementary Table 3).

In a secondary analysis, we included 1,642 CpGs from two manuscripts (6,9) fulfilling a FDR p-value<0.05. Among all the selected FDR results, 195 CpGs could not be analyzed in our study as they did not pass the quality control. We did not replicate any of the 1,447 CpGs analyzed (Table 2 and Supplementary Table 3). In the REGICOR study, the top hits were located in an intergenic region on chromosome 1 (cg10893043, p-value=6.79·10^−5^) and in the *PXK* and *ARSA* genes (cg16560256, p-value= 2.23·10^−04^; cg11953250, p-value=3.64·10^−04^) for PM_2.5_.

**Table 2:**
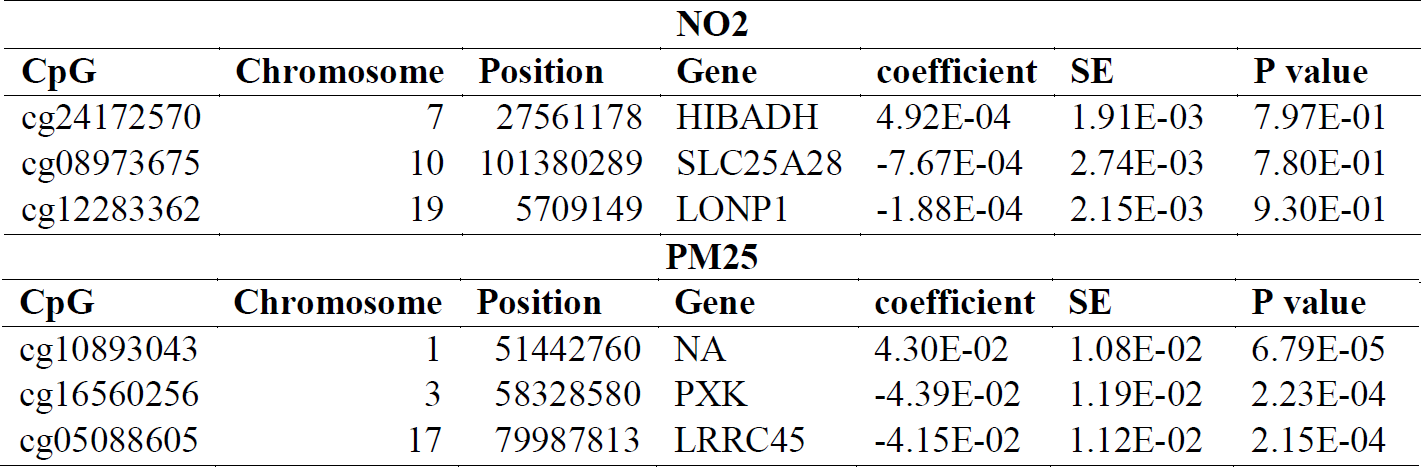
Top hits of the replication of previously published Cps associated with NO_2_ and PM_25_ exposures.

## DISCUSSION

This population-based and cross-sectional study did not identify new loci or replicate loci previously identified as showing differential methylation related to long-term exposure to air pollution.

The lack of positive results in our study should be interpreted with caution and some methodological issues must be considered. First, our study is underpowered to detect small effect size associations. The statistical power of our study was estimated (Supplementary Table 1 and 2) and is similar to previous studies. Second, we defined strict criteria to consider an association as statistically significant based on the Bonferroni multiple comparisons correction. This approach is more conservative than the FDR p-value used in other studies (6,9). However, in the replication effort we also included those CpGs that were identified using the FDR criteria and we did not replicate any of these CpGs in our sample. Third, the exposure to air pollution and its variability is lower than that observed in other studies, limiting our capability to identify real associations. Fourth, the exposure assessment was estimated using land use regression models. Although this methodology is commonly used and we followed the ESCAPE protocol, (18,19) using validated exposure estimations, (21) some exposure misclassification could still be present, reducing the power to detect the association of interest in our study. Fifth, the replication was carried in a case-control study that may have a different methylation pattern. Finally, we estimated long-term exposure to air pollution whereas other studies have analyzed the association between short-term exposure and DNA methylation; however, in studies that analyzed several time exposures, the longer the exposure the higher the number of loci showing differential methylation (9).

Despite these limitations that should be considered, we would highlight some of the results observed in this study. In the discovery effort, we identified 7 CpGs associated with PM_10_, 18 related to PM_2.5_, 6 with PM_coarse_, 28 with NO_X_, 32 with NO_2_, and 9 related to traffic at the nearest street in the REGICOR study. However, none of them were validated in the EPIC-Italy study. In our effort to replicate previous findings, we identified one locus located in an intergenic region on chromosome 1 (cg10893043, p-value=6.79·10-5) as potentially associated with PM_2.5_. The cg10893043 is close to the *CDKN2C* gene. The protein encoded by this gene is a member of the INK4 family of cyclin-dependent kinase inhibitors and regulates cell growth by controlling cell-cycle G1 progression (37). Some studies have shown that the expression of this gene inhibits the growth of human cells in animal models and have suggested a potential role in tumorigenesis (38).

Among the strengths of our study, we would mention the standardized methodology following the ESCAPE protocol that was consistently used to assess air pollution exposure. This methodology was validated in our population and was used to assess long-term air pollution exposure. Moreover, we applied a commonly used methodology to assess DNA methylation at the genome-wide level and a standardized methodology with both a discovery and an independent validation population.

## CONCLUSIONS

The results of our study are negative as we did not identify any new genomic loci associated with long-term air pollution and we did not replicate any previously identified loci. However, these negative results should be interpreted with caution. New joint efforts, increasing the statistical power of the analysis and the variability of the exposure to air pollution, and considering both short- and long-term exposure, are warranted to assess the potential association between air pollution and DNA methylation.

## LIST OF ABBREVIATIONS

WHO: World Health Organization.
REGICOR: REgistre GIroní del COR.
EPIC-Italy: The Italy center of European Prospective Investigation into Cancer and Nutrition.
450K: Infinium Human Methylation450 BeadChip.
CpG: Cytosine-phosphate-guanine.
PM_10_: Particulate matter with an aerodynamic diameter of <10μm.
PM_2.5_: Particulate matter with an aerodynamic diameter of <2.5μm.
PM_coarse_: The difference between PM_10_ and PM_2.5_.
NO_X_: Nitrogen oxides.
NO_2_: Nitrogen dioxide
LUR: Land use regression.
FDR: False Discovery Rate.

## DECLARATIONS

### Ethics approval and consent to participate

All participants in both studies (REGICOR and EPIC-Italy) signed an informed consent; the studies were approved by the local ethic committees (PSMAR CEIC-2012/4729/I) and followed national legislation and the Declaration of Helsinki criteria.

### Consent for publication

Not applicable.

### Availability of data and material

The datasets used and/or analyzed during the current study are available from the corresponding author [RE] on reasonable request.

### Authors’ contributions

- Conception or design of the study: SS-B, NK, JM, XB, RE

- Acquisition of data for the study: NK, JM, XB, RE

- Analysis of data for the manuscript: SS-B, IS, MP, XB

- Interpretation of data for the manuscript: SS-B, AF-S, AP, NK, JM, XB, RE

- Drafted the manuscript: SS-B, RE

- Revised the manuscript critically for important intellectual content: AF-S, AP, IS, MP, NK, JM, XB

- All the authors have approved the final version of the manuscript AND agree to be accountable for all aspects of the work in ensuring that questions related to the accuracy or integrity of any part of the work are appropriately investigated and resolved.

## Acknowledgments

We thank Elaine M. Lilly, PhD, for revising and editing the English text.

## REFERENCES

1. Organization WH. WHO: Air Pollution. 2018. http://www.who.int/airpollution/en/. Acccessed Feb 2018.

2. de F.C. Lichtenfels AJ, van der Plaat DA, de Jong K, van Diemen CC, Postma DS, Nedeljkovic I, et al. Long-term Air Pollution Exposure, Genome-wide DNA Methylation and Lung Function in the LifeLines Cohort Study. Environmental Health Perspectives. 2018;126:027004.

3. Plusquin M, Guida F, Polidoro S, Vermeulen R, Raaschou-Nielsen O, Campanella G, et al. DNA methylation and exposure to ambient air pollution in two prospective cohorts. Environment International. 2017;108:127–36.

4. Zhong J, Karlsson O, Wang G, Li J, Guo Y, Lin X, et al. B vitamins attenuate the epigenetic effects of ambient fine particles in a pilot human intervention trial. Proceedings of the National Academy of Sciences. 2017;114:3503–8.

5. Chi GC, Liu Y, MacDonald JW, Barr RG, Donohue KM, Hensley MD, et al. Long-term outdoor air pollution and DNA methylation in circulating monocytes: results from the Multi-Ethnic Study of Atherosclerosis (MESA). Environmental Health. 2016;15:119.

6. Gruzieva O, Xu CJ, Breton C V., Annesi-Maesano I, Antó JM, Auffray C, et al. Epigenome-wide meta-analysis of methylation in children related to prenatal NO2air pollution exposure. Environmental Health Perspectives. 2017;125:104–10.

7. Goodrich JM, Reddy P, Naidoo RN, Asharam K, Batterman S, Dolinoy DC. Prenatal exposures and DNA methylation in newborns: a pilot study in Durban, South Africa. Environ Sci: Processes Impacts. 2016;18:908–17.

8. Breton C, Gao L, Yao J, Siegmund K, Lurmann F, Gilliland F. Particulate matter, the newborn methylome, and cardio-respiratory health outcomes in childhood. Environ Epigenet. 2016;2:dvw005.

9. Panni T, Mehta AJ, Schwartz JD, Baccarelli AA, Just AC, Wolf K, et al. Genome-wide analysis of DNA methylation and fine particulate matter air pollution in three study populations: KORA F3, KORA F4, and the normative aging study. Environmental Health Perspectives. 2016;124:983–90.

10. Jiang R, Jones MJ, Sava F, Kobor MS, Carlsten C. Short-term diesel exhaust inhalation in a controlled human crossover study is associated with changes in DNA methylation of circulating mononuclear cells in asthmatics. Part Fibre Toxicol. 2014;11:71.

11. Grau M, Subirana I, Elosua R, Solanas P, Ramos R, Masiá R, et al. Trends in cardiovascular risk factor prevalence (1995-2000-2005) in northeastern Spain. European journal of cardiovascular prevention and rehabilitation?: official journal of the European Society of Cardiology, Working Groups on Epidemiology & Prevention and Cardiac Rehabilitation and Exercise Physiology. 2007; 653–9.

12. Beulens JWJ, Monninkhof EM, Monique Verschuren WM, van der Schouw YT, Smit J, Ocke MC, et al. Cohort profile: The EPIC-NL study. International Journal of Epidemiology. 2010;39:1170–8.

13. Palli D, Berrino F, Vineis P, Tumino R, Panico S, Masala G, et al. A molecular epidemiology project on diet and cancer: the EPIC-Italy Prospective Study. Design and baseline characteristics of participants. Tumori. 2003;89:586–93.

14. van Veldhoven K, Polidoro S, Baglietto L, Severi G, Sacerdote C, Panico S, et al. Epigenome-wide association study reveals decreased average methylation levels years before breast cancer diagnosis. Clinical Epigenetics. 2015;7:67.

15. Bibikova M, Barnes B, Tsan C, Ho V, Klotzle B, Le JM, et al. High density DNA methylation array with single CpG site resolution. Genomics. 2011; 98:288–95.

16. Sandoval J, Heyn HA, Moran S, Serra-Musach J, Pujana MA, Bibikova M, et al. Validation of a DNA methylation microarray for 450,000 CpG sites in the human genome. Epigenetics. 2011;6:692–702.

17. Sayols-Baixeras S, Subirana I, Lluis-Ganella C, Civeira F, Roquer J, Do A, et al. Identification and validation of seven new loci showing differential DNA methylation related to serum lipid profile: an epigenome-wide approach. The REGICOR study. Human Molecular Genetics. 2016;25:4556–65.

18. Beelen R, Hoek G, Vienneau D, Eeftens M, Dimakopoulou K, Pedeli X, et al. Development of NO2 and NOx land use regression models for estimating air pollution exposure in 36 study areas in Europe - The ESCAPE project. Atmospheric Environment. 2013;72:10–23.

19. Eeftens M, Beelen R, De Hoogh K, Bellander T, Cesaroni G, Cirach M, et al. Development of land use regression models for PM2.5, PM 2.5 absorbance, PM10 and PMcoarse in 20 European study areas; Results of the ESCAPE project. Environmental Science and Technology. 2012;46:11195–205.

20. Rivera M, Basagaña X, Aguilera I, Foraster M, Agis D, de Groot E, et al. Association between long-term exposure to traffic-related air pollution and subclinical atherosclerosis: The REGICOR study. Environmental Health Perspectives. 2013; 121:223–30.

21. Basagaña X, Aguilera I, Rivera M, Agis D, Foraster M, Marrugat J, et al. Measurement error in epidemiologic studies of air pollution based on land-use regression models. American Journal of Epidemiology. 2013;178:1342–6.

22. World Health Organisation. Manual of The MONICA Project. 2000. http://www.ktl.fi/publications/monica/manual/index.htm. Acccessed Feb 2018.

23. Baena-Díez JM, Alzamora-Sas MT, Grau M, Subirana I, Vila J, Torán P, et al. [Validity of the MONICA cardiovascular questionnaire compared with clinical records]. Gaceta sanitaria. 2010;23:519–25.

24. Houseman EA, Accomando WP, Koestler DC, Christensen BC, Marsit CJ, Nelson HH, et al. DNA methylation arrays as surrogate measures of cell mixture distribution. BMC Bioinformatics. 2012; 13:86.

25. Aryee MJ, Jaffe AE, Corrada-Bravo H, Ladd-Acosta C, Feinberg AP, Hansen KD, et al. Minfi: A flexible and comprehensive Bioconductor package for the analysis of Infinium DNA methylation microarrays. Bioinformatics. 2014;30(10):1363–9.

26. Jeffrey T. Leek and W. Evan Johnson and Hilary S. Parker and Andrew E. Jaffe and John D. Storey. sva: Surrogate Variable Analysis. R package version 3.10.0.

27. Jiang C-L, He S-W, Zhang Y-D, Duan H-X, Huang T, Huang Y-C, et al. Air pollution and DNA methylation alterations in lung cancer: A systematic and comparative study. Oncotarget. 2017;8:1369–91.

28. Anto JM, Bousquet J, Akdis M, Auffray C, Keil T, Momas I, et al. Mechanisms of the Development of Allergy (MeDALL): Introducing novel concepts in allergy phenotypes. Journal of Allergy and Clinical Immunology. 2017; 139:388–99.

29. Mostafavi N, Vlaanderen J, Portengen L, Chadeau-Hyam M, Modig L, Palli D, et al. Associations between genome-wide gene expression and ambient nitrogen oxides (NOx). Epidemiology. 2017; 28:320–8.

30. Gref A, Merid SK, Gruzieva O, Ballereau S, Becker A, Bellander T, et al. Genome-wide interaction analysis of air pollution exposure and childhood asthma with functional follow-up. American Journal of Respiratory and Critical Care Medicine. 2017;195:1373–83.

31. Somineni HK, Zhang X, Biagini Myers JM, Kovacic MB, Ulm A, Jurcak N, et al. Ten-eleven translocation 1 (TET1) methylation is associated with childhood asthma and traffic-related air pollution. Journal of Allergy and Clinical Immunology. 2016;137:797–805.

32. Hong X, Wang X. Epigenetics and Development of Food Allergy (FA) in Early Childhood. Current Allergy and Asthma Reports. 2014; 14:460.

33. Janssen BG, Godderis L, Pieters N, Poels K, Kiciński M, Cuypers A, et al. Placental DNA hypomethylation in association with particulate air pollution in early life. Particle and Fibre Toxicology. 2013;10:22.

34. Wang T, Garcia JG, Zhang W. Epigenetic Regulation in Particulate Matter-Mediated Cardiopulmonary Toxicities: A Systems Biology Perspective. Curr Pharmacogenomics Person Med. 2012;10:314–21.

35. Holloway JW, Savarimuthu Francis S, Fong KM, Yang I a. Genomics and the respiratory effects of air pollution exposure. Respirology. 2012;17:590–600.

36. Christensen BC, Marsit CJ. Epigenomics in environmental health. Frontiers in Genetics. 2011;2:84.

37. Cánepa ET, Scassa ME, Ceruti JM, Marazita MC, Carcagno AL, Sirkin PF, et al. INK4 proteins, a family of mammalian CDK inhibitors with novel biological functions. IUBMB Life. 2007; 59:419–26.

38. van Veelen W, Klompmaker R, Gloerich M, van Gasteren CJR, Kalkhoven E, Berger R, et al. P18 is a tumor suppressor gene involved in human medullary thyroid carcinoma and pheochromocytoma development. International journal of cancer. 2009;124:339–45.

